# Duo-activation of PKA and PKG by PDE1 inhibition facilitates proteasomal degradation of misfolded proteins and protects against proteinopathy

**DOI:** 10.1101/543785

**Authors:** Hanming Zhang, Bo Pan, Mark D. Rekhter, Alfred L. Goldberg, Xuejun Wang

**Affiliations:** Division of Basic Biomedical Sciences, University of South Dakota Sanford School of Medicine, Vermillion, SD 57069.; Lilly Research Laboratories, Lilly Corporate Center, Indianapolis, IN 46285.; Department of Cell Biology, Harvard Medical School, Boston, MA

**Keywords:** proteasome, cAMP-dependent protein kinase, cGMP-dependent protein kinase, phosphodiesterase 1 inhibitor, proteinopathy, heart failure with preserved ejection fraction

## Abstract

No current treatment is intended to target cardiac proteotoxicity or can reduce mortality of heart failure with preserved ejection fraction (HFpEF), a prevalent form of heart failure (HF). Selective degradation of misfolded proteins by the ubiquitin-proteasome system (UPS) is vital to the cell. Proteasome impairment is recently implicated in HF genesis. Activation of the cGMP-protein kinase G (PKG) or the cAMP-protein kinase A (PKA) pathways facilitates proteasome functioning. Phosphodiesterase 1 (PDE1) hydrolyzes both cyclic nucleotides and accounts for the majority of PDE activities in human myocardium. Here we report the preclinical therapeutic efficacy and a new mechanism of action of PDE1 inhibition (IC86430) for cardiac proteinopathy caused by Arg120Gly missense mutant αB-crystallin (CryAB^R120G^). In mice expressing GFPdgn, an inverse reporter of UPS proteolytic activity, IC86430 treatment increased myocardial 26S proteasome activities and substantially decreased GFPdgn protein levels. Myocardial PDE1A expression was highly upregulated in CryAB^R120G^ mice. HFpEF was detected in CryAB^R120G^ mice at 4 months; IC86430 treatment initiated at this stage markedly attenuated HFpEF, substantially delayed mouse premature death, increased myocardial levels of Ser14-phosphorylated Rpn6, and reduced the steady state level of the misfolded CryAB species in these mice. In cultured cardiomyocytes, IC86430 treatment increased proteasome activities and accelerated proteasomal degradation of GFPu and CryAB^R120G^ in a PKA- and PKG- dependent manner. We conclude that PDE1 inhibition induces PKA- and PKG-mediated promotion of proteasomal degradation of misfolded proteins in cardiomyocytes and effectively treats HFpEF caused by CryAB^R120G^; hence, PDE1 inhibition represents a potentially new therapeutic strategy for HFpEF and heart disease with increased proteotoxic stress.

**One Sentence Summary:** PDE1 inhibition enhances proteasomal degradation of misfolded proteins in a PKA and PKG dependent manner and protects against cardiac proteinopathy and heart failure with preserved ejection fraction.

## INTRODUCTION

Heart failure (HF) is a clinical syndrome in which the heart fails to pump enough blood to meet the body’s need for oxygen and nutrients. It can be broadly divided into HF with reduced ejection fraction (HFrEF) and HF with preserved ejection fraction (HFpEF). Current guidelines classify HF with EF < 40% as HFrEF. Conversely, patients with HFpEF, mainly characterized by diastolic malfunction, have a normal EF (>50%) (*1*). In the United States, approximately 50% of HF cases are HFpEF (*2*). Moreover, the prevalence of HFpEF relative to HFrEF has been increasing over time and HFpEF is projected to become the most common type of HF (*1*). Despite tremendous therapeutic advances in improving HFrEF mortality in the past decades, currently there is no effective pharmacological intervention available for HFpEF (*3*). Hence, HFpEF represents a large unmet need in cardiovascular medicine. Cardiac proteinopathy belongs to a family of cardiac diseases caused by increased expression of misfolded proteins and resultant proteotoxicity in cardiomyocytes (*4*). Emerging evidence suggests that increased proteotoxic stress (IPTS) plays an important role in the progression from a large subset of heart diseases to HF (*5*). As exemplified by desmin-related cardiomyopathy, restrictive cardiomyopathy or HFpEF is commonly seen in patients with cardiac proteinopathy (*6*). Similarly to HFpEF, no evidence-based clinical strategy is currently available for treating cardiac proteinopathy or for targeting cardiac proteotoxicity.

Targeted clearance of misfolded proteins by the ubiquitin-proteasome system (UPS) is pivotal to protein quality control (PQC) which acts to minimize the amount and toxicity of misfolded proteins in the cell (*7*). UPS-mediated proteolysis begins with covalent attachment of a chain of ubiquitin molecules to a substrate protein molecule via a highly coordinated cascade of enzymatic reactions, a process known as ubiquitination; the polyubiquitinated protein molecule is subsequently degraded by the 26S proteasome (*5, 8*). PQC inadequacy due to proteasome functional insufficiency (PFI) has been implicated in a large subset of human heart disease during their progression to HF and studies using animal models have established a major pathogenic role for PFI in cardiac proteinopathy, ischemia-reperfusion (I/R) injury, and diabetic cardiomyopathy (*10-16*). Hence, improving cardiac proteasome functioning is implicated as a potential therapeutic strategy for HF treatment. Moreover, recent advances in cell biology excitingly unravel the proteasome as a nodal point for controlling UPS proteolysis (*17, 18*). Our previous study has shown that cGMP-dependent protein kinase (PKG) stimulates proteasome proteolytic activities, improves cardiac UPS performance, and facilitates the degradation of misfolded proteins in cardiomyocytes (*19*). Additionally, cAMP-dependent protein kinase (PKA) was also shown to stimulate proteasome activity and accelerate proteasomal degradation of misfolded proteins in cultured cells through directly phosphorylating the 19S subunit RPN6/PSMD11 at Ser14 (*20*). Hence, it is conceivable, but remains to be formally tested, that duo-activation of PKG and PKA can facilitate proteasomal degradation of misfolded proteins in cardiomyocytes and thereby effectively treat heart disease with IPTS.

By breaking down cAMP and/or cGMP in a specific compartment of the cell, the cyclic nucleotide phosphodiesterase (PDE) plays a key role in cyclic nucleotide-mediated signaling amplitude, duration and compartmentalization in the cell. The PDE superfamily consists of 11 family members, PDE1 through PDE11 (*21*). PDE1s are duo-substrate PDEs and account for the majority PDE activity in human myocardium (*22*). The PDE1 family is encoded by three distinct genes, *PDE1A, 1B* and *1C*, with PDE1A and PDE1C expressed in the heart and vessels. PDE1A is mostly expressed in rodents and displays a higher affinity for cGMP than cAMP, whereas PDE1C is predominantly expressed in human myocardium and displays similar affinity to both cyclic nucleotides (*23, 24*). Several studies have shown an important role of PDE1 in cardiovascular regulation. A PDE1 selective inhibitor (IC86340) was shown to attenuate isoproterenol-induced cardiac hypertrophy and fibrosis in mice, with the mechanism involving stimulating the cGMP-PKG axis (*25, 26*). PDE1C depletion exhibits protective effect on pressure overload induced cardiac remodeling in a PKA-dependent manner (*27*). Given that both PKG and PKA positively regulate proteasome functioning, we sought to test the hypothesis that PDE1 inhibition augment proteasomal degradation of misfolded proteins and protect against cardiac proteinopathy, with the present study.

Here we report that myocardial PDE1A expression was substantially increased in a mouse model of cardiac IPTS, including at its HFpEF stage; PDE1 inhibition increased proteasome activities and the degradation of a known UPS substrate in the heart; and chronic treatment of CryAB^R120G^-based cardiac proteinopathy mice at their HFpEF stage with a PDE1-specific inhibitor (IC86430) improved cardiac diastolic function and remarkably delayed mouse premature death, which was associated with substantial increases in PKA-mediated proteasome phosphorylation and decreases in misfolded CryAB. In cultured cardiomyocytes, PDE1 inhibition increased 26S proteasome activities, enhanced degradation of a known UPS substrate in a PKA- and PKG- dependent manner and promoted proteasomal degradation of CryAB^R120G^, a misfolded protein known to cause human disease. Our data demonstrate that PDE1 inhibition, via activation of PKA and PKG, increases proteasome activities, promotes proteasomal degradation of misfolded proteins and protects against cardiac proteinopathy-based HFpEF, representing a potentially novel therapeutic strategy for HFpEF and heart disease with IPTS.

## RESULTS

### Upregulated myocardial PDE1A in a mouse model of cardiac proteinopathy

We first determined changes in myocardial expression of PDE1A in a well-established transgenic (tg) mouse model of cardiac proteinopathy induced by cardiomyocyte-restricted expression of CryAB^R120G^ (*28*), a misfolded protein known to cause human disease (*29, 30*). We examined non-tg (NTG), CryAB^R120G^ tg, wild-type CryAB (CryAB^WT^) tg mice at 4 months and 6 months of age when cardiac proteinopathy and cardiac malfunction are readily discernable in the CryAB^R120G^ tg mice. Western blot analyses with a validated anti-PDE1A antibody (Fig. S1) revealed that myocardial PDE1A protein levels were significantly increased in CryAB^R120G^ tg mice compared with NTG littermates (*p*=0.046 at 4 months, *p*=0.017 at 6 months, Fig.1, A and B). Myocardial PDE1A protein was not discernibly altered by overexpression of CryAB^WT^ (Fig. 1, C and D), suggesting that the upregulation of myocardial PDE1A is specific to the CryAB^R120G^ based-proteinopathy. Moreover, consistent with the protein data, RT-PCR analysis revealed that myocardial PDE1A mRNA levels were significantly increased in the CryAB^R120G^ mice compared to NTG littermates (*p*<0.05, Fig. 1, E and F), whereas no significant difference was detected between NTG and CryAB^WT^ tg mice (Fig. 1, E and F).

**Fig. 1.**
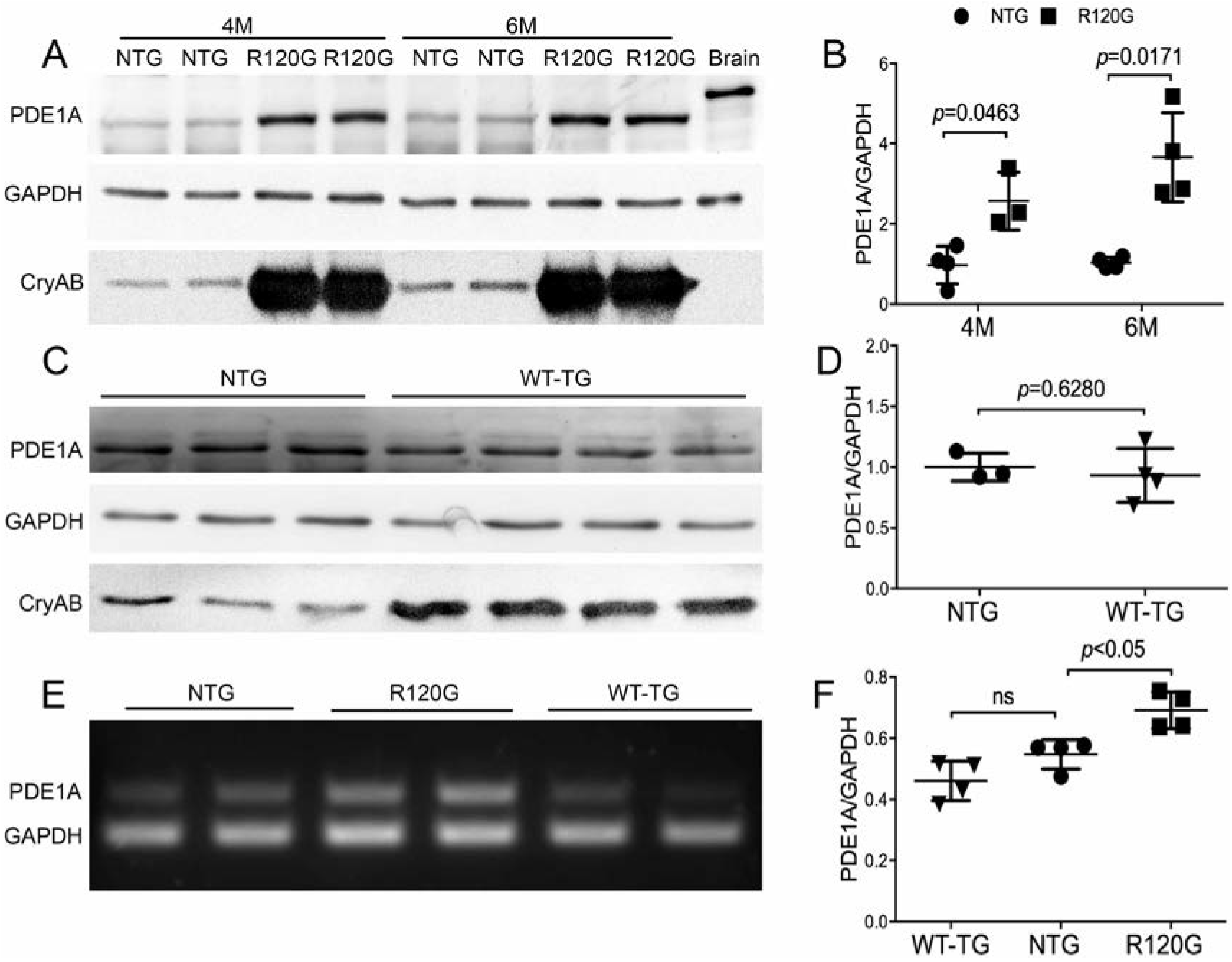
Myocardial PDE1A expression in mouse hearts overexpressing CryAB^R120G^ or wild type CryAB. Total protein and total RNA from ventricular myocardial samples were used for western blot analyses (**A∼D**) and RT-PCR (**E**, **F**), respectively. **A** and **B**, Representative images (**A**) and pooled densitometry data (**B**) of western blot analyses for PDE1A in 4-month or 6-month-old CryAB^R120G^ tg (R120G) and their non-tg (NTG) littermate mice. For each time point, 4 NTG (2 males + 2 males) and 4 R120G (2 males + 2 females) mice were used. **C** and **D**, western blot analysis for myocardial PDE1A in CryAB^WT^ tg (WT-TG) and NTG control mice at 6 months. Littermate NTG (1 male + 2 females) and WT-TG (2 males + 2 females) were used. **E** and **F**, Representative image (**E**) and pooled densitometry data (**F**) of RT-PCR analyses for myocardial PDE1A mRNA levels in NTG, R120G, and WT-TG mice at 6 months. For each genotype, 2 males and 2 females were tested. *p* Values are derived from two-tailed unpaired *t*-test with Welch’s correction (B, D) or one-way ANOVA followed by Tukey’s multiple comparison tests (F).

### Decreasing GFPdgn protein levels in mouse hearts by PDE1 inhibition

Activation of PKG promotes UPS proteolytic function (*19*). Cell culture studies carried out with non-cardiomyocyte cells have showed that cAMP/PKA also stimulates proteasome activities (*20, 31*). Since PDE1 hydrolyzes both cAMP and cGMP, we sought to determine whether PDE1 inhibition in mice is capable of enhancing myocardial UPS degradation of GFPdgn, a well-established UPS reporter substrate and a slightly shorter version of GFPu which is a green fluorescence protein (GFP) modified by carboxyl fusion of the degron CL1 (*32, 33*). We examined changes in myocardial GFPdgn expression at 6 hours after administration of IC86430 (3mg/kg, i.p.), a PDE1-specific inhibitor previously known as IC86340 (*26*). Myocardial GFPdgn protein levels were ∼40% lower in the IC86430-treated mice than that in vehicle-treated mice (*p*=0.041, Fig. 2, A and B), with the GFPdgn reduction evidenced primarily in the cardiomyocyte compartment (Fig. 2C); however, GFPdgn mRNA levels were comparable between the two groups (*p*=0.898, Fig. 2, D and E), indicating that the cardiac reduction of GFPdgn proteins by PDE1 inhibition occurs at the post transcriptional level. The 26S proteasome exists in two forms: doubly capped (i.e., a 20S core is capped at both ends by the 19S complex) and singly capped (i.e., only one end of the 20S core is capped by a 19S complex) (*34*). When the cytosolic fractions from the IC86430-treated GFPdgn mice were analyzed with native gel electrophoresis followed by an in-gel proteasome peptidase activity assay, they showed increased peptidase activities both in singly (*p*=0.0006 vs. vehicle, Fig. 2, F and G) and doubly capped species (*p*=0.0026 vs. vehicle, Fig. 2, F and G), whereas proteasome assembly was not discernibly altered (Fig. 2, F and H). Taken together, these results suggest that PDE1 inhibition by IC86430 increases myocardial 26S proteasome activities, stimulates UPS proteolytic function, and facilitates the clearance of a surrogate misfolded protein in vivo.

**Fig. 2.**
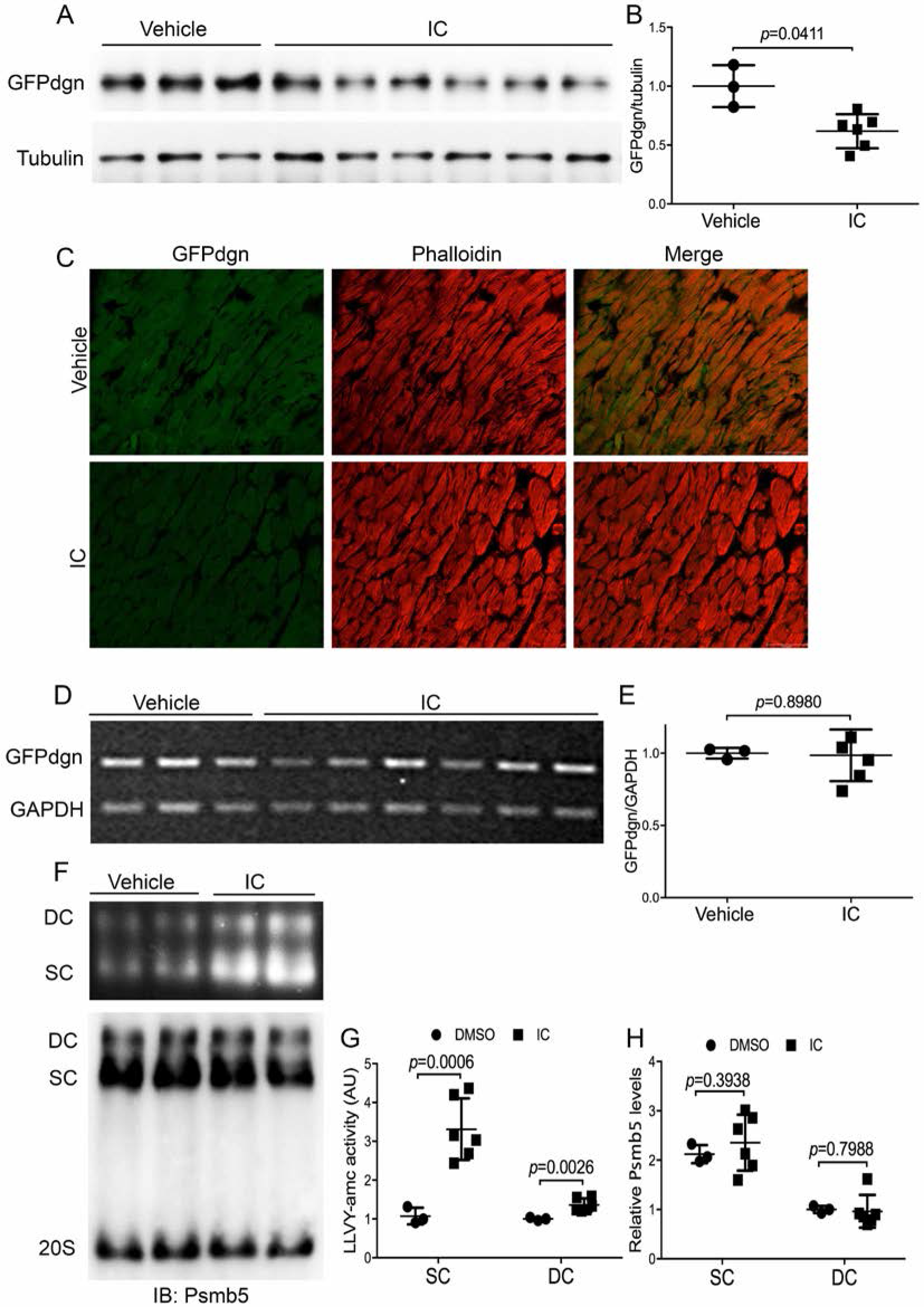
Effects of IC86430 on GFPdgn expression and the assembly and activity of 26S proteasomes in mouse hearts. Adult GFPdgn tg mice were treated with IC86430 (IC, 3mg/kg, i.p., n=6) or equivalent amount of vehicle (n=3). Ventricular myocardium was sampled 6 h after injection for the indicated assays. **A** and **B**, western blot analysis images (**A**) and pooled densitometry data (**B**) for the indicated proteins. **C**, Representative fluorescence confocal micrographs of IC or vehicle treated GFPdgn mouse ventricular myocardium. **D** and **E**, RT-PCR image (**D**) and densitometry data (**E**) of myocardial GFPdgn mRNA levels. Duplex RT-PCR for GFPdgn and GAPDH were performed. **F**∼**H**, Analyses for the activity and abundance of 26S proteasomes after native gel electrophoresis. Myocardial crude protein extracts were subject to native gel electrophoresis and then in-gel peptidase activity assays using suc-LLVY-amc as the fluorogenic substrate; the fractionated proteins in the native gel were then transferred to a PVDF membrane and subject to immunoblot for Psmb5 to assess the abundance of the doubly capped 26S proteasomes (DC) and singly capped 26S proteasomes (SC). Shown are representative images of the in-gel peptidase activity assay (upper **F**) and corresponding western blot (lower **F**) as well as the pooled densitometry data (**G**, **H**) of respective assays. All *p* values in this figure were derived from two-tailed unpaired *t*-test with Welch’s correction.

### Protection against HFpEF and premature death of CryAB^R120G^ mice by chronic PDE1 inhibition

Since PDE1 inhibition can enhance cardiac UPS proteolytic function and PFI has proven to be a major pathogenic factor in cardiac proteinopathy (*11*), we decided to test whether chronic PDE1 inhibition is able to mitigate CryAB^R120G^-based cardiac proteinopathy in mice. Line 134 CryAB^R120G^ tg mice develop well-characterized cardiac proteinopathy (*28*). In brief, the animals do not show any cardiac pathology at 1 month of age but they do show restrictive cardiomyopathy at 3 months and heart failure at 6 months and die between 6 and 7 months of age (*28*). Hence, this model serves as an excellent cardiac proteinopathy model for studying pathogenesis and experimental intervention. We started the treatment at exactly 4 months of age when cardiac hypertrophy and heart malfunction are readily discernible (Fig. 3) but the mice are approximately 2 months younger than their median lifespan (*28, 35*). A cohort of line 134 CryAB^R120G^ tg mice were randomly assigned to either IC86430 (3mg/kg/day; TG-IC) or vehicle control (equivalent amount of 60% DMSO in saline, TG-Vehicle) treatment groups and the treatment was designed to last for 4 weeks via osmotic mini-pumps. We carried out Kaplan-Meier survival analysis, in addition to serial echocardiography performed immediately before (baseline) and 4-, 6-, and 8- weeks after mini-pump implantation. As expected, no statistically significant difference in any of the echocardiographic parameters was detected between TG-IC and TG-Vehicle groups before initiation of the treatment (Table S1). However, data from echocardiography performed at the day immediately before mini-pump implantation reveal a moderate HFpEF phenotype in the CryAB^R120G^ tg mice, as reflected by significant decreases in left ventricular (LV) end diastolic volume (LVEDV), in stroke volume (SV), and in cardiac output (CO), as well as increases in end diastolic LV posterior wall thickness (LVPWd) along with unchanged ejection fraction (EF), compared with the NTG littermate mice (Fig. 3). Importantly, these echocardiographic abnormities were substantially attenuated (CO, LVPWd) or even normalized to the NTG control levels (LVEDV, SV), without altering EF and the heart rate, at the end of 4 week-treatment in the TG-IC group compared with the TG-Vehicle group (Fig. S2 and Fig. 4, A, C, E, G, and I). Although virtually all echocardiographic parameters became indistinguishable between the TG-IC and TG-Vehicle groups by 4 weeks after termination of the treatment (i.e., 8 weeks after initiation of treatment), analyses of the combined data of the entire 8 weeks of assessment period using the area under curve (AUC) of each parameter as the index convincingly show significantly improvement in LVEDV, SV, and CO in the TG-IC group compared with the TG-Vehicle group while EF and heart rate remained comparable between the two groups throughout (Fig. 4, B, D, F and J). Taken together, these data demonstrate that the IC86430 treatment improves cardiac function and effectively attenuates HFpEF in the CryAB^R120G^ tg mice.

**Fig. 3.**
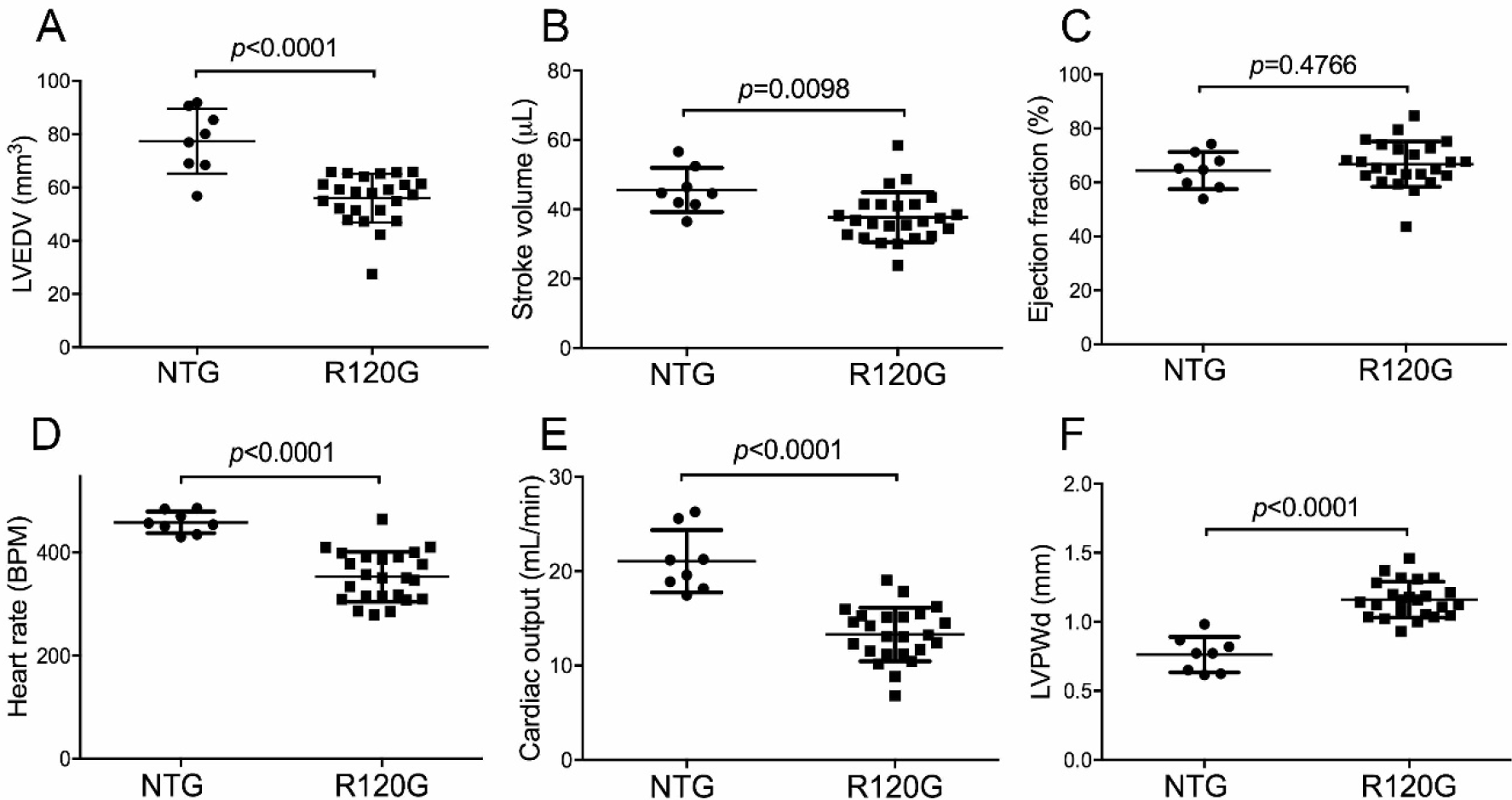
HFpEF is detected in CryAB^R120G^ tg mice at 4 months of age. Parameters shown are derived from echocardiography on CryAB^R120G^ tg (R120G) and sex-matched NTG littermate mice at exactly 4 months of age, i.e., the day immediately before the mini-pump implantation. LVEDV, left ventricular (LV) end-diastolic volume; LVPWd, end-diastolic LV posterior wall thickness. Littermate NTG (4 males + 4 females) and R120G (10 males + 14 females) were used. Two-tailed unpaired *t*-test.

**Fig. 4.**
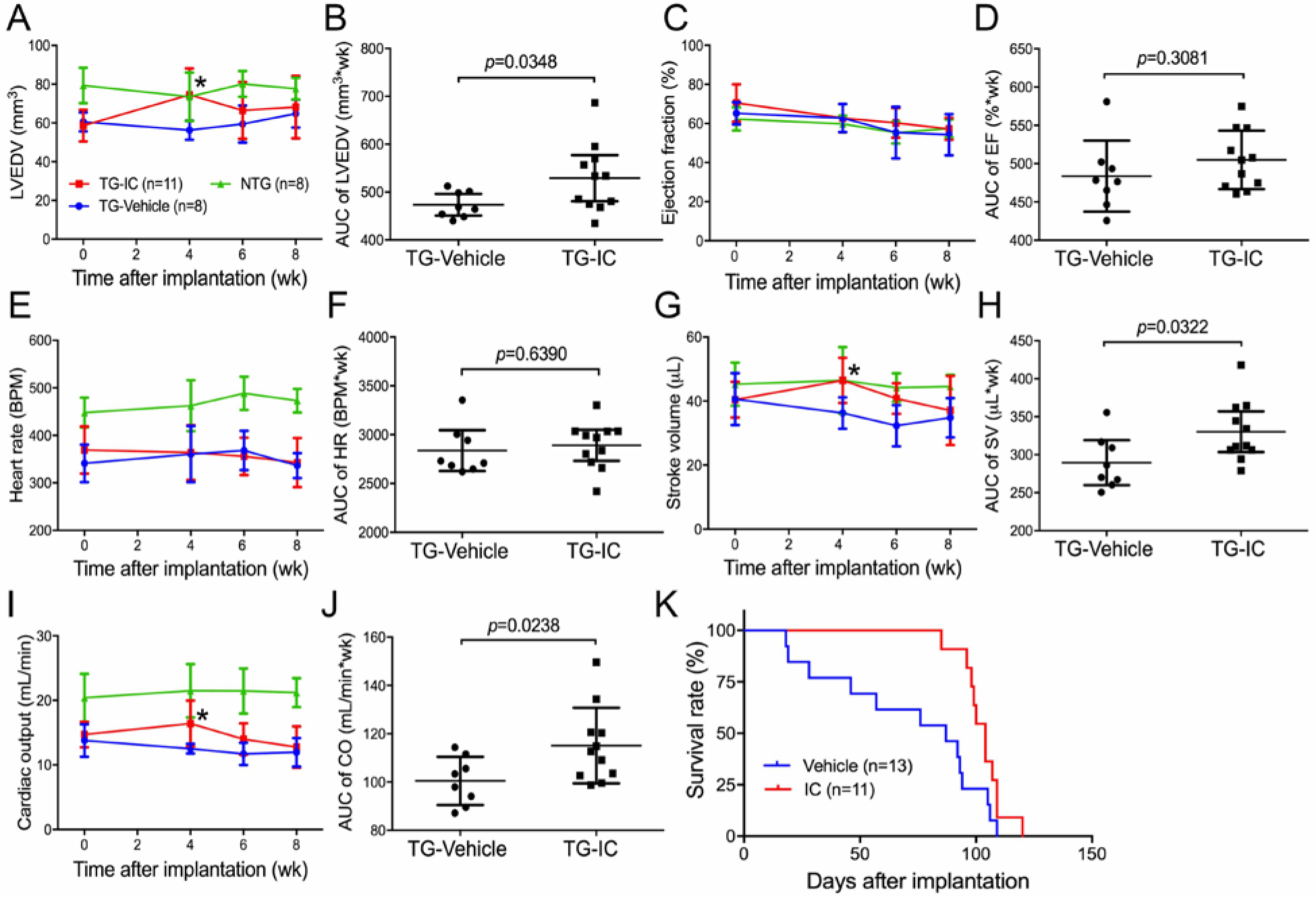
Effects of PDE1 inhibition on CryAB^R120G^ tg mouse left ventricular (LV) function and morphometry and lifespan. The treatment of CryAB^R120G^ tg mice with IC86430 (3mg/kg/day; TG-IC) or vehicle control (TG-Vehicle) was initiated at exactly 4 months of age and lasted for 4 weeks via subcutaneous implantation of mini-osmotic pumps. The mice were observed daily for survival and subject to serial echocardiography on the day before (baseline) as well as 4, 6, and 8 weeks after the mini-pump implantation. **A - J**, LV morphometry and function parameters derived from the serial echocardiography of the TG-IC (4 males + 7 females) or TG-Vehicle (4 males + 4 females) mice or of age-matched non-tg littermates (NTG; 4 males +4 females) without any intervention. The stacked line chart of each panel summarizes the time course of changes in the indicated parameter. Mean ± SD; \**p*<0.05 TG-IC vs. TG-Vehicle of the same time point (two-way repeated measures ANOVA followed by Bonferroni’s test). The scatter dot plot of each panel presents the area under curve (AUC) of the indicated parameter versus time obtained using the trapezoidal rule. Each dot represents a mouse; mean ± SD are superimposed; two-tailed unpaired *t*-test with Welch’s correction were used. **K**, Kaplan-Meier survival analysis. The median lifespans of the TG-IC tg group (4 males + 7 females) and the TG-Vehicle (6 males + 7 females) groups are 224 days and 207 days, respectively. *p*=0.0235, log-rank test.

Consistent with the echocardiography data, Kaplan-Meier survival analysis of the cohort revealed that IC86430 treatment remarkably delayed the premature death of the CryAB^R120G^ tg mice (*p*= 0.0235, Fig. 4K). Notably, the TG-Vehicle group displayed a survival curve that is comparable to that was previously reported for this line of tg mice (*11, 36*). Importantly, during the 4-week IC86430 treatment and within the 6 weeks following termination of the IC86430 treatment, none of the TG-IC mice died but ∼50% of the TG-Vehicle mice died during this period (Fig. 4K), demonstrating a striking and long-lasting therapeutic benefit from the IC86430 treatment.

### Decreasing oligomeric CryAB^R120G^ and increasing Ser14-phopshorylated Rpn6 in proteinopathic mouse hearts by PDE1 inhibition

Given that IC86430 treatment enhances degradation of a surrogate UPS substrate in mouse hearts (Fig. 2A), we next asked whether PDE1 inhibition would promote proteasomal degradation of misfolded CryAB^R120G^ proteins in the heart. A recent study developed a novel method to distinguish proteasomal degradation of misfolding-prone proteins from other cellular mechanisms that regulate total protein level (*37*). In this assay, total protein lysate is separated into soluble (supernatant, NS) and insoluble fractions (pellet, NI) using a NP-40 lysis buffer. The pellet is then suspended in an SDS buffer and is further divided into SDS soluble (SS) and SDS insoluble fractions (SR). The SS fraction was rapidly increased by proteasome inhibition, suggesting that it derives from a misfolded species and can be readily prevented by the proteasome; therefore, changes in the level of a stably expressed misfolding-prone protein in the SS fraction inversely reflect proteasome proteolytic activity (*37*). Hence, we examined the SS fraction of CryAB at the end of the 4-week treatment. As expected, the protein level of the SS fraction of CryAB was barely detectable in NTG mouse hearts but was significantly increased in the CryAB^R120G^ tg mice (Fig. S3); importantly, the increase was remarkably attenuated by IC86430 treatment (*p*=0.0014; Fig. 5, A and B). Furthermore, the protein level of Ser14-phosphorylated RPN6 (p-Rpn6), which has proven to be increased by PKA activation (*20*), was significantly increased in the TG-IC mice compared with TG-Vehicle mice (*p*=0.0189; Fig. 5, C and D), indicating that myocardial PKA-mediated proteasome stimulation is enhanced by PDE1 inhibition. Taken together, these in vivo data highly support that PDE1 inhibition leads to PKA-mediated phosphorylation of Rpn6 and thereby increases proteasome activity and facilitates proteasomal degradation of misfolded proteins in the heart.

**Fig. 5.**
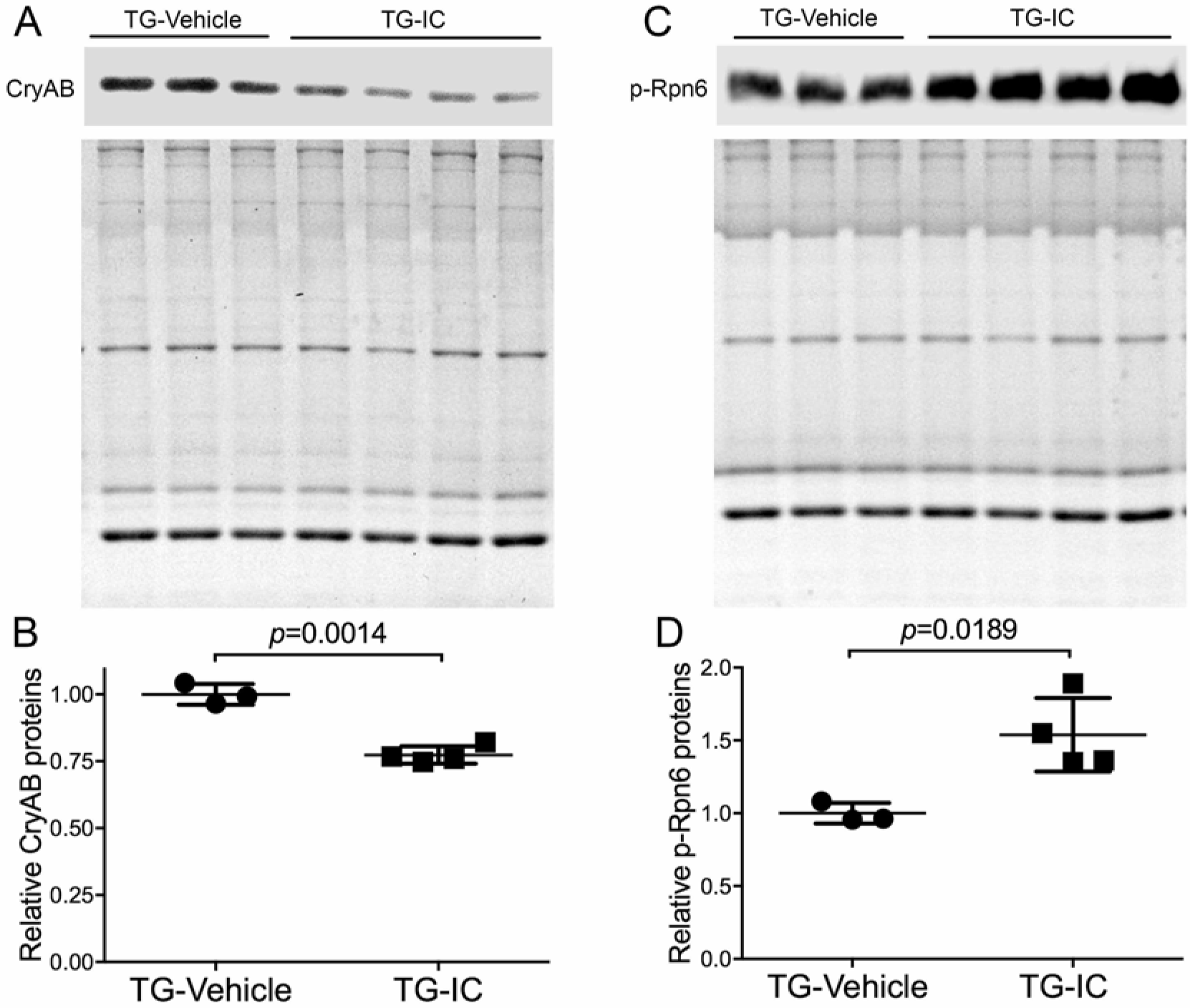
Effects of IC86430 treatment on myocardial abundance of oligomeric CryAB and Ser14-phosphorylated Rpn6 in CryAB^R1220G^ tg mice. A separate cohort of CryAB^R120G^ mice subject to the same treatment as described in **Figure 4** were sacrificed immediately at the end of the 4-week treatment and ventricular myocardium was sampled for protein analyses. **A** and **B**, the western blot images (**A**) and summary of densitometry data (**B**) of the NP-40 insoluble but SDS soluble fraction of CryAB; **C** and **D**, Representative images (**C**) and pooled densitometry data (**D**) of western blot analyses for Ser14-phosphorylated Rpn6 (p-Rpn6) are shown. The stain-free total protein imaging technology was utilized to visualize all proteins in the SDS-PAGE gel used for the western blot analysis. The total protein signals of corresponding lane of the gel (the bottom image of panels **A** and **C**) are used as the in-lane loading control. Each lane (**A**, **C**) or each dot (**B**, **D**) represents an independent mouse. The difference between the TG-IC (3 males + 1 female) and the TG-Vehicle (2 males +1 female) group is evaluated statistically using two-tailed unpaired *t*-test with Welch’s correction.

### Proteasome enhancement by PDE1 inhibition in cultured cardiomyocytes

Next we tested the effect of PDE1 inhibition on proteasome proteolytic activities in cultured cardiomyocytes using both small peptide substrates and a full length protein surrogate. In-gel proteasome peptidase activity assays using the cytosolic fractions revealed significantly higher peptidase activities in both singly capped and doubly capped proteasomes from the IC86430-treated NRVMs than those from the vehicle treated cells (Fig. 6, A to C). To determine whether PDE1 inhibition enhances the degradation of a known substrate of the UPS in a cardiomyocyte-autonomous fashion, we tested the effect of IC86430 treatment on the stability of GFPu in cultured NRVMs. Adenoviral vectors expressing GFPu (Ad-GFPu) or a red fluorescent protein (Ad-RFP) were used. The RFP has a much longer half-life than GFPu and is not an efficient UPS substrate whereas the GFPu is a proven UPS substrate (*19*). The Ad-RFP has exactly the same expression control elements for RFP as the Ad-GFPu for GFPu; hence, RFP protein levels can serve as a valid control for both viral infection efficiency and protein synthesis when a mixture of Ad-RFP and Ad-GFPu is used (*19, 38, 39*). In Ad-GFPu and Ad-RFP co-infected NRVMs, treatment with IC86430 dose-dependently decreased the ratio of GFPu to RFP, indicative of increased degradation of GFPu (Fig. 6, D and E). Moreover, the cycloheximide (CHX) chase assay showed that PDE1 inhibition significantly shortened the half-life of GFPu (*p*=0.0026, Figure. 6, F and H), demonstrating that PDE1 inhibition promotes UPS proteolytic function in the cultured cardiomyocytes. Taken together, these results strongly suggest that UPS proteolytic function enhancement by PDE1 inhibition is at least in part attributable to enhanced activities of the 26S proteasome.

**Fig. 6.**
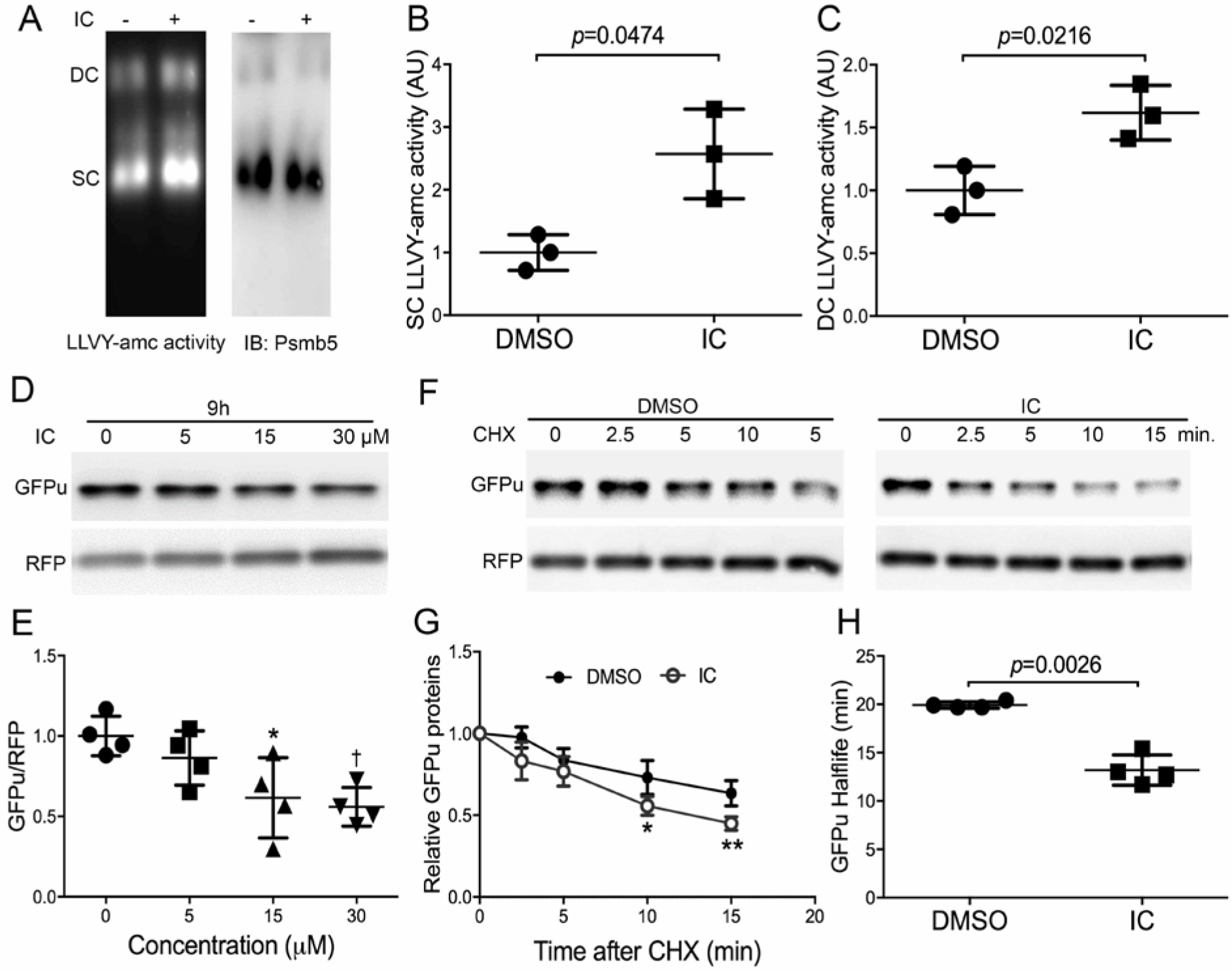
PDE1 inhibition by IC86430 increases proteasomal peptidase activities and promotes proteasomal degradation of GFPu in cultured neonate rat ventricular myocytes (NRVMs). **A∼C**, in-gel proteasome peptidase activity assays followed by immunoblotting (IB) for Psmb5 were performed as described in Figure 2F. NRVMs in culture were treated with IC86430 (IC, 15 μM) for 9 hours before being harvested for crude protein extraction. A fraction of each sample was used for the in-gel proteasomal peptidase assay (**A**∼**C**). Representative images (**A**) and pooled data of 3 biological repeats (**B, C**) of the cleavage of the LLVY-AMC fluorogenic substrate are shown. Two-tailed unpaired t-test with Welch’s correction. **D** and **E**, representative images (**D**) and pooled densitometry data (**E**) of western blot analyses for the steady state protein levels of the co-overexpressed GFPu and RFP. Cultured NRVMs were co-infected with a mixture of recombinant adenoviruses expressing GFPu and RFP in serum-free medium for 6 h. Cells were then cultured in DMEM containing 2% fetal bovine serum for 48 hours before IC treatment. Total cell lysates collected after 9 hours of IC treatment at the indicated dosages were analyzed. \**p*<0.05, †*p*<0.01 vs. the 0 μM group; n=4 repeats; one way ANOVA followed by Dunnett’s test. **F∼H**, cycloheximide (CHX) chase assays for GFPu degradation. CHX treatment (100 μM) was started 5 hours after the treatment of IC (15 μM) or DMSO. In each run of the CHX chase, the GFPu chase was started 15 min after addition of CHX to the culture dish and the GFPu image density at this time point is set as 1 arbitrary unit (AU) and the GFPu levels of subsequent time points were calculated relative to it. Representative western blot images (**F**), summarized GFPu decay curves (G), and the GFPu half-lives derived repeats (**H**) are shown. \**p*<0.05, \*\**p* <0.01vs. DMSO group; two-tailed unpaired t-test with Welch’s correction.

### Accelerating proteasomal degradation of CryAB^R120G^ by PDE1 inhibition

To determine whether the PDE1 inhibition-induced acceleration of misfolded protein degradation in the heart is cardiomyocyte-autonomous, we sought to further examine the effect of PDE1 inhibition on degradation of a human disease-linked misfolded protein, CryAB^R120G^ in cultured NRVMs. HA-tagged CryAB^R120G^ was overexpressed in NRVMs via adenoviral gene delivery. Consistent with in vivo data, PDE1 inhibition by IC86430 significantly reduced CryAB^R120G^ in the NP40-insoluble but SDS-soluble fraction (*p*=0.0099, Fig. 7, A and B). The reduction of CryAB^R120G^ protein level by IC86430 is proteasome-dependent as the reduction was reversed in the presence of proteasome inhibitor bortezomib (BZM, Fig. 7, C and D). Furthermore, the CHX chase assay revealed that PDE1 inhibition significantly shortened CryAB^R120G^ protein half-life (*p*=0.0197, Fig 7, E and G). Together, these data further confirm that PDE1 inhibition enhances proteasome-mediated degradation of misfolded proteins in cardiomyocytes.

**Fig. 7.**
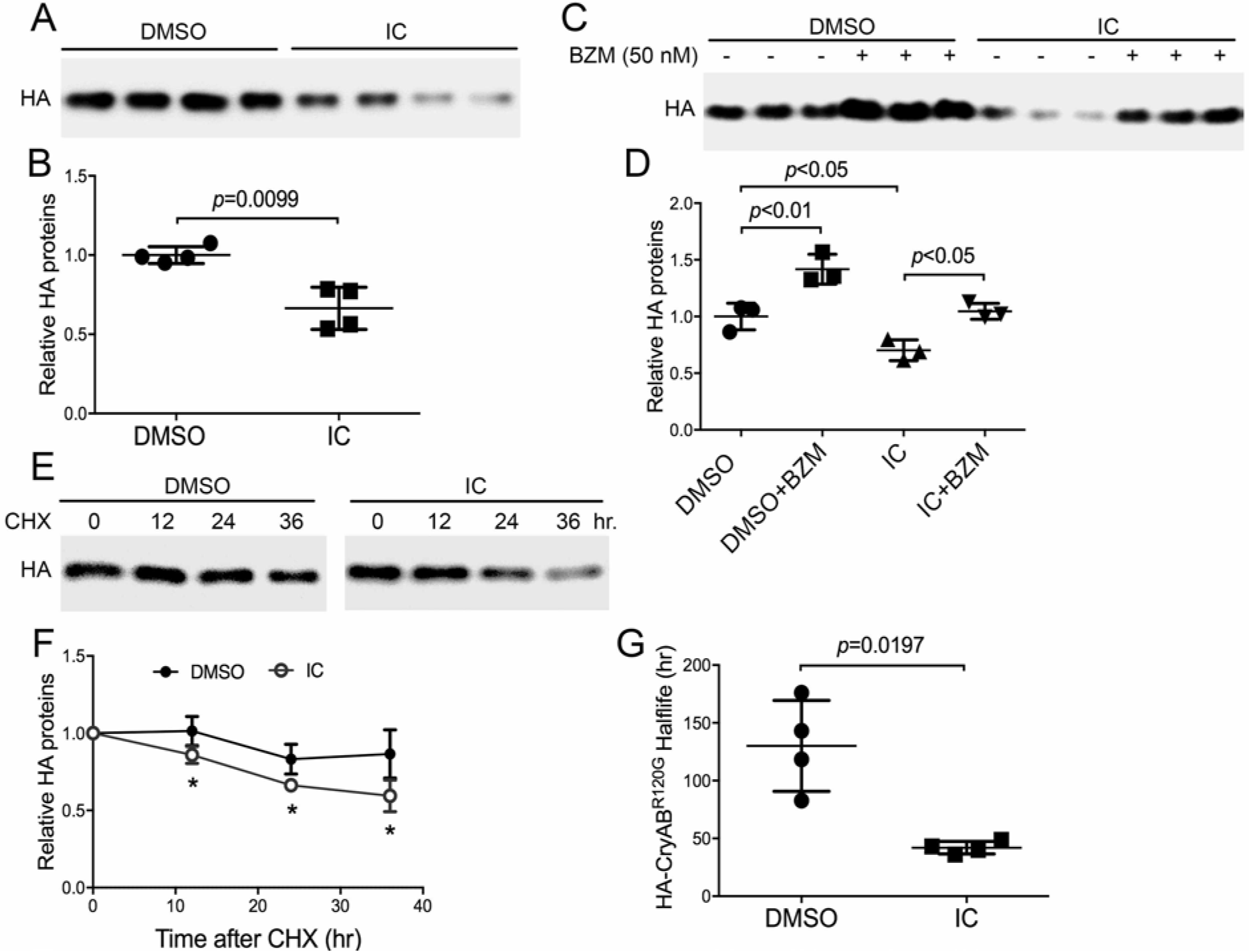
PDE1 inhibition enhances proteasomal degradation of CrAB^R120G^, a bona fide misfolded protein. Cultured NRVMs were infected with adenoviruses expressing HA-tagged CryAB^R120G^ (Ad-HA-CryAB^R120G^) in serum-free medium for 6 hours. Cells were then cultured in DMEM containing 2% serum for 24 hours before treated with IC86430 (IC, 30 μM) or vehicle control (DMSO). **A** and **B**, Representative image (**A**) and pooled densitometry data (**B**) of western blot analyses for HA-CryAB^R120G^ in the NP40 insoluble but SDS soluble fraction of NRVMs treated with IC for 6 h. **C** and **D**, Reduction of a misfolded species of CryAB level by IC is proteasome dependent. The proteasome inhibitor bortezomib (BZM, 50 nM) or volume corrected DMSO was applied to NRVM cultures 10 h after the initiation of IC or DMSO treatment. Six hours later, the cells were harvested for extraction of the NP-40 insoluble but SDS-soluble fraction of proteins. Representative images (**C**) and the pooled densitometry data from three repeats (**D**) of western blot analyses for HA-CryAB^R120G^ are shown. Two-way ANOVA followed by Tukey’s multiple comparisons test, n=3 repeats. **E**∼**G**, CHX chase assays of HA-CryAB^R120G^. CHX (50 μM) treatment was started after 16h of IC or DMSO treatment. Total protein extracts from NRVMs harvested at the indicated time points after CHX administration were subject to western blotting for HA-CryAB^R120G^. Representative images (**E**), HA-CryAB^R120G^ decay curves (**F**), and the derived HA-CryAB^R120G^ half-lives (**G**) are presented. \**p*<0.05 vs. DMSO group, two-tailed unpaired t-test with Welch’s correction; n=4 repeats.

### Attenuation of PDE1 inhibition’s proteasomal enhancement by inhibiting PKA or PKG

Experiments performed with non-cardiac cells have demonstrated that stimulating the cAMP/PKA pathway increases 26S proteasome activities and promotes proteasomal degradation of several disease-linked misfolded proteins through phosphorylating RPN6 at its Ser14 (*20*). To test whether cAMP/PKA would do the same in cardiomyocytes, we treated cultured NRVMs with forskolin, a potent activator of adenylate cyclase, to increase cAMP, in the absence or the presence of a PKA inhibitor H89. Western blot analyses revealed that forskolin remarkably increased the level of Ser14-phosphorylated RPN6 and this increase was completely blocked by H89 co-treatment (Fig. 8A). CHX chase assays further showed forskolin significantly shortened the half-life of GFPu in the cultured NRVMs (Fig. 8, B and C), confirming that the proteasome phosphoregulation by cAMP/PKA does occur in cardiomyocytes. Previously we have reported that cGMP/PKG positively regulates 26S proteasome activities and the proteasomal degradation of misfolded proteins in cardiomyocytes (*19*). Since PDE1 breaks down both cAMP and cGMP, we then sought to test the role of PKA or PKG activation in the acceleration of GFPu degradation by PDE1 inhibition. Both PKG inhibitor (KT5823) and PKA inhibitor (H89) significantly attenuated the reduction of GFPu by IC86430 treatment (Fig. 8, D to G), indicating that both PKA and PKG activation mediate PDE1 inhibition-induced proteasomal function enhancement in cardiomyocytes.

**Fig. 8.**
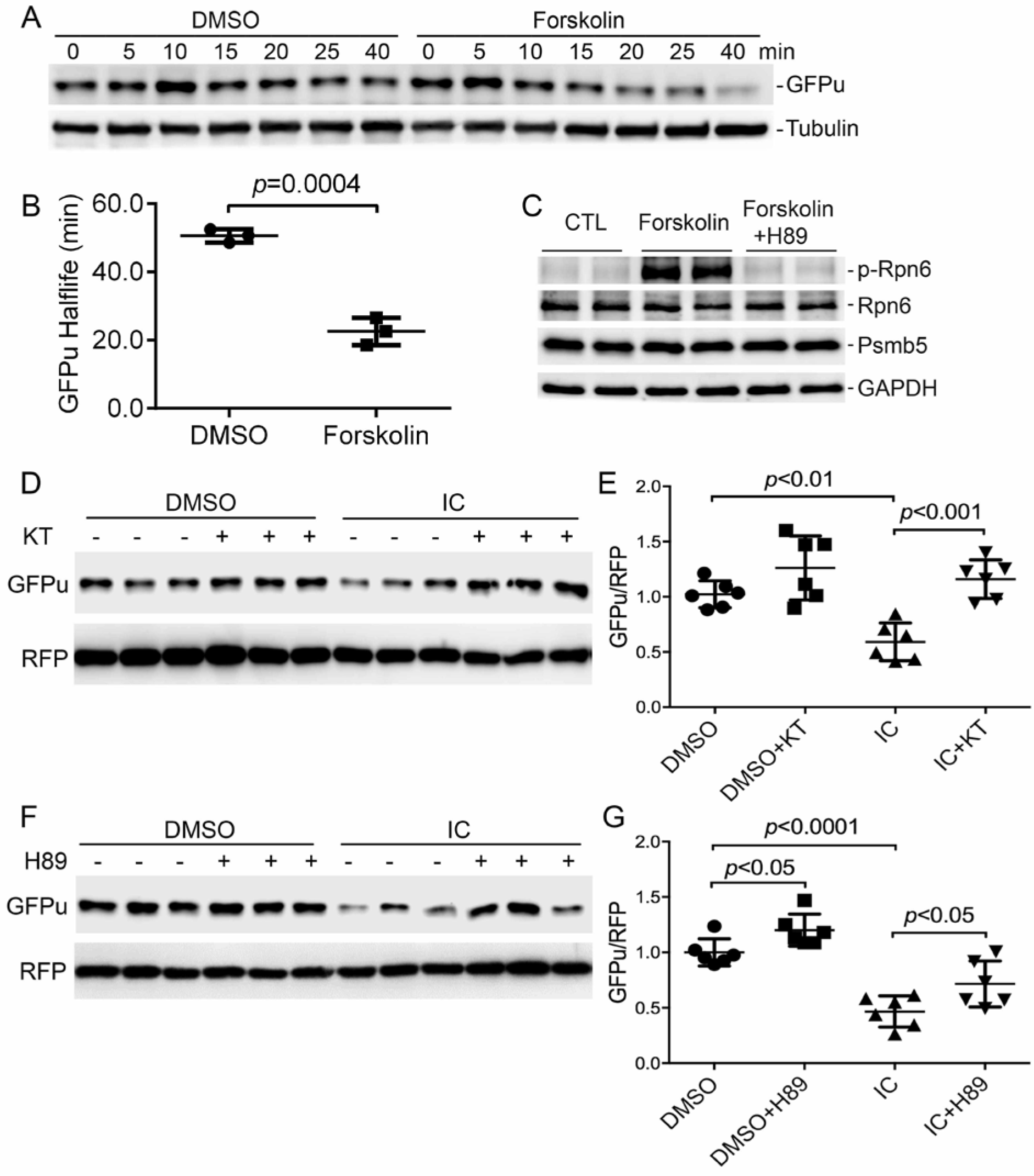
Accelerated degradation of GFPu by PDE1 inhibition in cardiomyocytes is PKA and PKG dependent. **A** and **B**, CHX chase assay for changes in GFPu protein stability induced by increasing cAMP in cardiomyocytes. NRVMs were infected with Ad-GFPu in serum-free medium for 2 h; the medium was then switched to 10% FBS medium for the reminder of the experiment. Forty-eight hours later, the treatment with forskolin (10 μM), a potent adenylyl cyclase activator, or with the vehicle control (DMSO) was initiated, which was 2 h before blocking protein synthesis with CHX (100 mg/ml). Cell lysates harvested at the indicated time point post-CHX were subject to SDS-PAGE and western blot analysis for GFPu and β-tubulin. Example western bot images (**A**) and the estimated GFPu protein half-lives (**B**) are shown. Two-tailed unpaired *t*-test with Welch’s correction. **C**, Western blot analyses for p-Rpn6 and other indicated proteins in NRVMs subject to the indicated treatment for 6 h. The PKA inhibitor H89 (5μM) was added 5 min before forskolin (10μM). **D**∼**G**, Western blot analyses for GFPu and RFP co-expressed in cardiomyocytes subject to the indicated treatments. Cultured NRVMs were co-infected with Ad-GFPu and Ad-RFP in serum free medium for 6 hours. The cells were then cultured in DMEM containing 2% serum for 24 hours before treatment indicated below. The treatment with a PKG inhibitor KT5823 (KT, 1 μM) or volume corrected vehicle (DMSO) was started 3 hours before IC86430 (IC) treatment (30 μM). The cells were harvested 9 hours later (**D**, **E**). For the PKA experiment, NRVMs were pre-treated with IC (30 μM) or volume corrected vehicle DMSO for 3 hours before H89 (5 μM) or volume corrected vehicle DMSO was administered. The cells were harvested 6 hours later (**F**, **G**). Representative western blot analyses (**D**, **F**) for GFPu with RFP probed as the loading control and a summary of the pooled densitometry (**E**, **G**) are shown. Two-way ANOVA followed by Tukey’s multiple comparisons test, n=6 repeats per manipulation.

## DISCUSSION

Prior to the present study, Miller *et al.* (*26*) demonstrated that PDE1A was upregulated in hypertrophied mouse hearts and isolated cardiomyocytes exposed to hypertrophic stimuli such as isoproterenol (ISO) or Angiotensin II (Ang II). Both pharmacological inhibition of PDE1 with IC86340 and gene silencing of PDE1A attenuated cardiac hypertrophy, an effect associated with activation of the cGMP-PKG axis (*26*). PDE1C was reported to play a critical role in pressure overload induced pathological cardiac remodeling and dysfunction in a PKA dependent manner (*27*). In addition, PDE1 inhibition with IC86340, as shown by Zhang et al., protected against doxorubicin-induced cardiac toxicity and dysfunction in vivo (*16*). Recently, emerging evidence showed that patients in the end-state HF display increased protein level of PDE1A (*15*) and increased mRNA level of PDE1C (*15, 27, 40*). In the present study, we used both in vitro and in vivo approaches to investigate the therapeutic effects and underlying mechanism of PDE1 inhibition on HFpEF induced by proteotoxicity and have gained the following novel findings: (1) PDE1A expression at both mRNA and protein levels is upregulated in mouse hearts with advanced cardiac proteinopathy induced by CryAB^R120G^, a human misfolded protein; (2) CryAB^R120G^ mice develop HFpEF at 4 months of age and chronic PDE1 inhibition initiated at this stage effectively blocks the progression of HFpEF and delays mouse premature death in this model; (3) PDE1 inhibition, likely acting on the cAMP-PKA axis as well as the cGMP-PKG pathway, promotes proteasomal degradation of misfolded proteins, thereby protecting cardiomyocytes from proteotoxicity. Taken together, we demonstrate here that duo-activation of PKA and PKG by PDE1 inhibition facilitates proteasomal degradation of misfolded proteins and thereby protects against cardiac diastolic malfunction caused by IPTS.

### Potential mechanisms underlying the protection of PDE1 inhibition against proteinopathy

The mechanisms underlying the beneficial effects of PDE1 inhibition on cardiac function are likely quite complex, given the potentially multi-pronged involvement of PDE1 (a dual PDE) in cardiac regulation. As elaborated below, findings of this study strongly support the notion that improving cardiac PQC via enhancing proteasomal degradation of misfolded proteins represents an important molecular mechanism underlying the therapeutic effect of PDE1 inhibition against HFpEF due to αB-crystallin mutation.

Firstly, our data compellingly support that PDE1 inhibition-induced UPS enhancement protects the heart from proteotoxic stress. The degradation of individual misfolded proteins is primarily performed by the UPS. When UPS proteolytic function is impaired or becomes inadequate, misfolded proteins undergo aberrant protein aggregation with the formation of oligomers in the initial step. The intermediate oligomers are believed to be toxic to the cell (*41*). In the present study, we showed that the NP40 insoluble but SDS soluble fraction of CryAB^R120G^ (i.e., the oligomers formed by misfolded CryAB) was significantly decreased by IC86430 treatment (Fig. 5, A and B). Additionally, PDE1 inhibition also reduced the oligomeric CryAB^R120G^ in cultured NRVMs (Fig. 7, A and B). The reduction of CryAB^R120G^ oligomers by PDE1 inhibition in cardiomyocytes is proteasome-dependent because the reduction was prevented by proteasome inhibition (Fig. 7C). Thus, the improvement of cardiac function by IC86430 treatment in the proteinopathic mice is at least in part attributable to the enhanced clearance of misfolded CryAB^R120G^ by the UPS. Importantly, accompanied with the reduction of misfolded proteins, the myocardial protein level of Ser14-phosphorylated Rpn6 (p-Rpn6) was markedly upregulated in the IC86430-treated CryAB^R120G^ tg mice (Fig. 5, C and D). The 26S proteasome consists of a 20S proteolytic core particle (the 20S) and a 19S regulatory particle (the 19S) attached at one or both ends of the 20S. The 19S is composed of at least 17 subunits, including regulatory particle non-ATPase (RPN) 1 to 12 and regulatory particle triple-A^+^ ATPase (RPT) 1 to 6 (*42*). Recent studies revealed that phosphorylation of Rpn6 at Ser14 by PKA increased proteasome activity and thereby enhanced proteasomal degradation of misfolded proteins in the tested cells which are non-cardiac cells (*20*); here we were able to confirm that PKA activation increases p-Rpn6 and facilitates degradation of a UPS surrogate substrate (GFPu) in cardiomyocytes as well (Fig. 8, A to C). Furthermore, in cultured NRVMs, we found that 26S proteasome peptidase activities were substantially increased (Fig. 6, A to C) and the degradation of GFPu was remarkably accelerated (Fig. 6, D to H) by PDE1 inhibition. The accelerated GFPu degradation by IC86430 was discernibly attenuated by inhibition of either PKA or PKG (Fig. 8, D to G), indicating that improving UPS functioning in cardiomyocytes by PDE1 inhibition depends on activation of both PKA and PKG, which is also consistent with PDE1 being a duo-substrate PDE. Our previous study has shown that PKG activation enhances proteasome peptidase activity and promotes degradation of misfolded proteins (*19*). Hence, it is very likely that PDE1 inhibition enhances both cAMP-PKA and cGMP-PKG dependent activation of the proteasome, thereby promoting proteasomal degradation of misfolded proteins in the heart.

Second, PFI is a major pathogenic factor for the model we used and correction of PFI is known to be cardioprotective. Enhancing cardiac proteasome function by cardiomyocyte-restricted overexpression of proteasome activator 28α (PA28α) remarkably reduces aberrant protein aggregation, slows down the progression of proteinopathy, and delays mouse premature death in this CryAB^R120G^-based proteinopathy mouse model and protects against acute myocardial ischemia-reperfusion injury, demonstrating that PFI is a major pathogenic factor in heart disease with IPTS (*11*). In agreement with findings of the present study, increasing cGMP and thereby activating PKG by PDE5 inhibition with sildenafil also has been demonstrated to increase myocardial proteasome activities, decrease myocardial CryAB^R120G^ protein levels, and slow down the disease progression in CryAB^R120G^ mice (*19*), the first in vivo demonstration that proteasome function can be pharmacologically enhanced to treat disease. The activation of PKA by PDE4 inhibition has been shown to protect against a mouse model of Alzheimer’s disease (*43*). No reported study has tested directly the effect of PKA activation on cardiac proteotoxicity but pharmacologically induced PKA activation was shown to increase myocardial proteasome assembly and proteasome activities (*44*). Therefore, there is compelling evidence that through PKA- and PKG-mediated proteasomal activation, PDE1 inhibition with IC86430 facilitates degradation of misfolded proteins in cardiomyocytes and thereby reduces cardiac proteotoxicity and protects the heart against IPTS. Although the cardiac proteasome impairment hypothesis had not been proposed until years after neural scientists hypothesized a pathogenic role for UPS impairment in neurodegeneration, cardiac experimentalists were able to take the lead to establish both genetically and pharmacologically that impaired proteasome function plays a major role in cardiac pathogenesis (*11, 12, 14, 19, 45*), even before colleague of neuroscience did so to neural disorders (*43, 46*). Importantly, pathological studies have revealed that accumulation of ubiquitinated proteins and pre-amyloid oligomers and reduction of proteasome activities are prevalent in explanted failing human hearts (*9, 47-49*), further supporting a major pathogenic role for proteasome impairment in a large subset of heart failure in humans (*41, 42, 50*). Therefore, findings of the present study provide a new compelling rationale for PDE1 inhibition to treat heart failure.

### Potential contributions of proteasome enhancement to most reported pharmacological benefits of PDE1 inhibition

One of the beneficial effects of PDE1 inhibition, as demonstrated by Miller *et al*. (*26*), is protection against pathological cardiac hypertrophy. They found PDE1 inhibition by IC86340 and PDE1A downregulation by shRNA attenuated phenylephrine (PE) induced hypertrophic responses in isolated rat cardiomyocytes in a PKG dependent manner, highly suggesting a PDE1A-cGMP/PKG axis in regulating cardiac hypertrophy in cardiomyocytes. However, the underlying mechanism by which PDE1A-mediated cGMP-PKG regulates cardiac hypertrophy remains elusive. More recently, Knight *et al*. reported that global *PDE1C* knockout significantly attenuated transverse aortic constriction (TAC)-induced cardiac remodeling and dysfunction, characterized by decreases in chamber dilation, myocardial hypertrophy, and interstitial fibrosis in mice (*27*). In line with the reported anti-hypertrophic effect of PDE1 inhibition (*25, 26*), we also observed a statistically significant decrease of posterior wall thickness at the end of 4-week IC86430 treatment in CryAB^R120G^ mice (Table S2). Pathological cardiac hypertrophy is a maladaptive response to increased stress and requires increased protein synthesis which inevitably increases production of misfolded proteins and proteotoxic stress (*7*). This is because (i) it has been reported that approximately one third of the newly synthesized polypeptides never became mature proteins and instead were co-translationally degraded by the UPS (*51*) and (ii) a hypertrophied cardiomyocyte demands for a greater PQC capacity to maintain the quality of the mature proteins. Hence, for quality control purposes, the demand for UPS-mediated proteolysis is increased in response to cardiac hypotrophy (*41*). In fact, UPS inadequacy and PFI have been observed in pressure overload cardiomyopathy and PFI activates the calcineurin-NFAT pathway (*45, 52-54*), an important signaling pathway to pathological hypertrophy. PQC improvement by removal of misfolded proteins and protein aggregates has been shown to attenuate cardiac hypertrophy (*11, 36*). Proteasome enhancement was also shown to protect against pressure overload induced right ventricular failure (*45*). Therefore, it is conceivable that PQC improvement resulting from PKA- and PKG- mediated proteasome enhancement contribute to the anti-hypertrophic effect of PDE1 inhibition.

Reduction of cardiomyocyte death is another reported benefit from PDE1 inhibition. PDE1C depletion or PDE1 inhibition using IC86340 abolished angiotensin II (AngII) or isoproterenol (ISO)-induced cell death in isolated cardiomyocytes. The anti-apoptotic effects of PDE1C deficiency or inhibition were mediated by the cAMP-PKA axis (*27*). Zhang, *et al*. further demonstrated that PDE1C co-localized with adenosine A2 receptors (A2Rs) in cardiomyocyte membrane and T-tubules and the anti-apoptotic effect of PDE1 inhibition is dependent on cAMP-generating A2Rs in cardiomyocytes, suggesting that PDE1C selectively regulates A2R-cAMP signaling in cardiomyocytes (*16*). Since impaired PQC resulting from PFI is sufficient to cause cardiomyocytes to die, proteasome enhancement may also contribute to the pro-survival effect of PDE1 inhibition. Overexpression of a polyglutamine oligomer or CryAB^R120G^ results in cardiomyocyte death and, conversely, accelerating the degradation of misfolded proteins improves cardiomyocyte survival (*19, 55*). Proteasome inhibition by MG132 alone is sufficient to induce cell death in cultured NRVMs (*56*). Beyond PQC, proteasomal degradation also positively regulates cell signaling pathways that support cardiomyocyte survival. Akt-mediated pro-survival signaling was impaired in a mouse genetic model of cardiomyocyte-restricted proteasomal inhibition during myocardial ischemia/reperfusion (I/R) injury, likely due to impairment of proteasome-mediated degradation of PTEN, rendering cardiomyocytes more vulnerable to apoptosis in response to ischemia/reperfusion (*13*).

In addition to acting on cardiomyocytes, PDE1 inhibition and PDE1A downregulation attenuated AngII- or transforming growth factor β (TGFβ)-induced cardiac myofibroblast activation and extracellular matrix (ECM) synthesis in cultures (*25*). The anti-fibrotic effect is dependent on the suppression of both cAMP/Epac1/Rap1 signaling and cGMP/PKG signaling (*25*). Interestingly, PDE1C, which is only expressed in cardiomyocytes and undetectable in cardiac fibroblasts, also regulates TGFβ-induced fibroblast activation, likely through a paracrine-dependent mechanism (*27*). Hence, both in vitro and in vivo evidence compellingly support an anti-fibrosis effect for PDE1 inhibition. It is well-known that myocardial fibrosis can be induced by loss of cardiomyocytes (*57*), known as replacement fibrosis; hence, the anti-fibrotic effect could be secondary to the protection against cell death by PDE1 inhibition. Even with reactive fibrosis where a paracrine mechanism may be more relevant, proteasome enhancement also could potentially participate in PDE1 inhibition’s suppressive effect, probably because proteasome enhancement may curtail TGFβ signaling; both TGFβ receptors and intracellular signaling mediators Smad2/3 are degraded by the UPS and, conversely, proteasome inhibition was shown to promote the TGFβ signaling (*58-60*).

### PDE1 inhibition as a potential strategy to treat HFpEF

Human patients with HFpEF often have both cardiac (e.g., defective ventricular filling, concentric hypertrophy, reduced SV and CO and preserved EF) and non-cardiac manifestations (e.g., obesity, insulin-resistance, and type II diabetes) and an integrative molecular basis for these manifestations remains murky. Despite the prevalence of HFpEF in humans, there is an apparent shortage of animal models that can completely mimic human HFpEF conditions, hindering the pathogenic and therapeutic exploration of HFpEF. Prior characterization of the CryAB^R120G^ tg mouse line used here revealed that these mice display no discernible abnormal cardiac phenotype at 1 month of age, develop concentric cardiac hypertrophy and diastolic malfunction at 3 months, and die of HF between 6 and 7 months (*28, 35*). In the present study, IC86430 treatment of CryAB^R120G^ mice was initiated at the disease stage when the animal displays all cardiac characteristics of moderate HFpEF as reflected by decreases in LVEDV, SV, CO, and LVIDd, increased LVPWd, as well as normal EF (Fig. 3 and Table S1). During the entire treatment session as well as the follow-up echocardiographic surveillance, the EF in the CryAB^R120G^ mice remained comparable to that in NTG mice (Fig. 4C) whereas the cardiac morphometric and functional changes characteristic of HFpEF became more pronounced in the TG-Vehicle group, indicating that the CryAB^R120G^ model is primarily an HFpEF model, at least during the treatment period. In fact, human cardiac proteinopathies, including those caused by the CryAB^R120G^ mutation, often present in the clinic as restrictive cardiomyopathy (*61-63*), a cousin if not an etiology of HFpEF. Importantly, we have demonstrated that a defined period of PDE1 inhibition with IC86430 can effectively diminish the HFpEF-associated cardiac morphological and functional deficits, including normalizing LVEDV and SV (Fig.4, A and G) and thereby improving CO (Fig. 4I), resulting in a significant delay in the premature death of the diseased mice (Fig. 4K). This is an extremely exciting discovery as HFpEF lacks an established medical treatment.

Recently, Hashimoto *et al*. reported that acute PDE1 inhibition using a compound (ITI-214) different from what we used here exerted cardioprotective effect on HF dogs induced by tachypacing (*15*). Interestingly, they observed acute PDE1 inhibition exerts positive inotropic, lusitropic, chronotropic, and arterial vasodilatory effects in mammalian hearts expressing primarily PDE1C and those effects are likely derived from modulation of cAMP; however, they were unable to detect the inotropic effects, though a mild systemic vasodilation and increase in heart rate, in normal mice where the dominant isoform is PDE1A. We did not test the acute effect of IC86430 treatment on cardiac mechanical performance in normal mice but we observed cardiac proteasome functional enhancement by a single dose of IC86430 in GFPdgn mice (Fig. 2), suggesting that IC86430 and ITI-214 may differ in their targeting preference of PED1 isoforms. By contrast, we report here that chronic PDE1 inhibition with IC86430 improves cardiac diastolic function and animal survival in a well-tested mouse model of cardiac proteotoxicity likely through enhanced proteasomal degradation of misfolded proteins in a cAMP- and cGMP-dependent manner. This seeming discrepancy may arise from multiple aspects of difference between the two studies. First of all, different PDE1 inhibitors were used; different PDE1 inhibitors might have preference in not only PDE1 isoforms but also subcellular compartments as implicated by inhibitors of other PDE families (*64*). It is known that PDE1A and PDE1C regulate cardiac structural remodeling and function likely with distinct mechanistic actions (*65*). Our pan-PDE1 inhibitor (IC86430) likely inhibits both PDE1A and PDE1C while ITI-214 is claimed as PDE1C-selective; therefore, it is not surprising that they yield different therapeutic effects. Second, the duration of treatment differs between the studies; mice were treated continuously for 4 weeks in our study whereas only acute effects were determined by Hashimoto, *et al*. (*15*). It should be pointed out that both our in vivo and in vitro experiments reveal that PDE1 inhibition by IC86430 can acutely enhance proteasome proteolytic function in murine cardiomyocytes as well, suggesting that difference in targeting preference of PDE1 isoforms between the two PDE1 inhibitors is likely the primary cause.

Notably, the IC86430 treatment in our survival study was initiated at an overt disease stage when aberrant protein aggregates, cardiac hypertrophy, and diastolic malfunction are discernible. The Kaplan-Meier survival results are striking, indicating the drug’s overall therapeutic efficacy is very encouraging regardless of its underlying mechanism. A couple of clinically translatable therapeutic approaches, including exercise and doxycycline treatment (*36, 66*), have been shown to improve the survival of the proteinopathy mouse model used in the present study and their survival benefit seemed to be more impressive than PDE1 inhibition by IC86430 treatment. In both prior reports, the intervention started earlier (at 1 month of age for the exercise experiment and 16 weeks of age for Doxycycline treatment) and both treatments lasted for the entire remaining lifetime of the mice treated; the IC86430 treatment in the present study, however, lasted for only 4 weeks. Thus, the observed lifespan elongation effects are not comparable. It remains to be tested but very likely that the survival improvement by IC86430 would be much greater should the treatment be started earlier or last longer.

In summary, we have demonstrated in the present study that PDE1 inhibition improves PQC and protects against proteotoxicity in an animal model of heart failure. Taken together, our data as well as other existing reports provide compelling evidence that PDE1 inhibition shall be explored as a new therapeutic strategy for HFpEF and heart disease with IPTS.

## MATERIALS AND METHODS

### Animal models

The NIH Guide for the Care and Use of Laboratory Animals was followed throughout the study. Animal protocols used in this study were approved by the University of South Dakota Institutional Animal Care and Use Committee. The creation and characterization of the transgenic (tg) mouse model expressing GFPdgn were reported before (*32*). GFPdgn is a slightly shorter version of GFPu. GFPu is an enhanced green fluorescence protein (GFP) that is modified by carboxyl fusion of degron CL1 (*67*). Both GFPu and GFPdgn are surrogates for misfolded proteins and well-proven UPS substrates (*32*). The tg mice with cardiomyocyte-restricted overexpression of CryAB^R120G^ or CryAB^WT^ were described before (*28*). CryAB^R120G^, CryAB^WT^, and GFPdgn tg mice were maintained in the FVB/N inbred background. Transgenic mice were identified by PCR analysis of genomic DNA isolated from toe or tail clips.

### Mouse PDE1 inhibitor treatment

A PDE1-selective inhibitor IC86430 was provided by Eli Lily and Company (Indianapolis, Indiana 46285, USA). Age- and sex-matched GFPdgn mice were treated with IC86430 (3mg/kg, i.p.) or equivalent amount of vehicle control (DMSO). At 6 hours after the injection, ventricular myocardium was collected for total RNA isolation and protein extraction for further analyses. Treatment to cohorts of age- and sex-matched line 134 CryAB^R120G^ tg mice with IC86430 (3 mg/kg/day) or vehicle control was initiated at exactly 4 months of age and lasted for 4 weeks using subcutaneous mini-osmotic pumps (Alzet 2004 model). Dose determination was based on previous reports (*25, 26*).

### Echocardiography

Trans-thoracic echocardiography was performed on mice using the VisualSonics Vevo 2100 system and a 30-MHZ probe as previously described (*68*).

### Neonatal rat ventricular myocyte (NRVM) culture and adenoviral infection

NRVMs were isolated from the ventricles of 2-day old Sprague-Dawley rats using the Cellutron Neomyocytes isolation system (Cellutron Life Technology, Baltimore, MD), plated on 6-cm plates at a density of 2.0 × 10^6^ cells in 10% FBS, and cultured as described before (*69*). The plated cells were then infected with adenoviruses harboring the expression cassette for GFPu (Ad-GFPu), red fluorescent protein (RFP) (Ad-RFP), or HA-tagged CryAB^R120G^ (Ad-HA-CryAB^R120G^) (*19*).

### Extraction of the NP-40 soluble and insoluble fractions

NRVMs were infected with Ad-HA-CryAB^R120G^ in serum free DMEM media for 6 hours. The cells were then cultured in DMEM containing 2% serum for 24 hours before protein extraction. Cells were washed in cold phosphate buffered saline (PBS) at pH 7.4, harvested into cell lysis buffer (50 mM Tris, pH 8.8, 100 mM NaCl, 5 mM MgCl_2_, 0.5% NP-40, 2 mM DTT) containing 1× complete protease and phosphatase cocktail (T-2496, A.G. Scientific, Inc.), and incubated on ice for 30 min. Cell lysates were then centrifuged at 17,000 × g for 15 min at 4 °C. The supernatant was collected as the NP-40-soluble fraction (NS). The pellet containing NP-40-insoluble fractions (NI) was re-suspended in cell pellet buffer (20 mM Tris, pH 8.0, 15 mM MgCl2, 2 mM DTT, 1x complete protease and phosphatase cocktail), followed by 30 min incubation on ice. The NI fractions were further solubilized in 3× SDS boiling buffer (6% SDS, 20 mM Tris, pH 8.0, 150 mM DTT) and boiled for 5 min. The NI fractions were then subject to SDS-PAGE and western blot analysis for the misfolded species of CryAB.

### Western blot analysis

Total proteins were extracted from ventricular myocardium or cultured NRVMs with 1× sampling buffer (41mM Tris-HCl, 1.2% SDS, 8% glycerol). Protein concentration was determined with bicinchoninic acid (BCA) reagents (Pierce biotechnology, Rockford, IL). Equal amounts of proteins were loaded to each lane of the SDS-polyacrylamide gel and fractionated via electrophoresis, transferred to a PVDF membrane using a Trans-blot apparatus (Bio-Rad, Hercules, CA) and immunodetected with anti-PDE1A (SC-50480, Santa Cruz biotechnology, 1:800), anti-GAPDH (G8795, Sigma-Aldrich; 1:1000), anti-HA (#3724, Cell Signaling; 1:1000), anti-GFP (SC-9996, Santa Cruz biotechnology, 1:1000), anti-tubulin (E7-s, DSHB, 1:1000), anti-CryAB (ADI-SPA-222, Enzo Life Sciences), or anti-Psmb5 (custom made (*13*), 1:1000) as the primary antibodies and appropriate horse radish peroxidase-conjugated secondary antibodies. The bound secondary antibodies were detected using the enhanced chemiluminescence (ECL) detection reagents (GE Healthcare, Piscataway, NJ). Blots were imaged and quantified using the Quantity-One^TM^ or Image Lab^TM^ software (Bio-Rad). For some of the blots, the total protein content derived from the stain-free protein imaging technology was used as in-lane loading control (*15, 16*).

### Cycloheximide (CHX) chase assay

To determine protein half-life of GFPu or HA-tagged CryAB^R120G^, NRVMs were cultured in DMEM in the presence of cycloheximide (50 μM; Sigma-Aldrich), which was used to block further protein synthesis. Cells were collected at various time points after administration of CHX and total proteins were extracted for western blot analysis for GFPu or HA-tagged CryAB^R120G^.

### RNA isolation and reverse transcription-polymerase chain reaction (RT-PCR)

Total RNA was extracted from ventricular myocardium using the Tri-Reagent (Molecular Research Center, Inc., Cincinnati, OH) as described before (*19*). The concentration of RNA was determined using Agilent RNA 6000 Nano assay (Agilent technologies, Inc. Germany) following the manufacturer’s instruction. For reverse transcription (RT) reaction, 1 μg of RNA was used as a template to generate cDNA using the SuperScript III First-Strand Synthesis kit (Invitrogen) and the RT was performed by following the manufacturer’s instructions. For the duplex PCR of a gene-of-interest and GAPDH, 2 μl of solution from the RT reaction and specific primers towards the target gene and GAPDH were used. The mRNA levels of the gene-of-interest were quantified by PCR at the minimum number of cycles (15 cycles) capable of detecting the PCR products within the linear amplification range. Mouse PDE1A, GFPdgn, and GAPDH were measured using the following primer pairs: PDE1A: 5’-CTAAAGATGAAC T GGAGGGATCTTCG GAAC-3’ (Forward) and 5’-TGGAGAAAATGGAAGCCCTAATTCAGC-3’ (Reverse); GFPdgn: 5’-TCTATATCATGGCCGACAAGCAGA-3’ (Forward) and 5’-ACTGGGTGCTC AGGTAGT GGTTGT-3’ (Reverse); GAPDH: 5’-ATGACATCAAGAAGGTGGTG-3’ (Forward) and 5’-CATACCA GGAAATGA GCTTG-3’ (Reverse).

### Native gel electrophoresis and in-gel proteasome peptidase activity assays

In-gel proteasome peptidase activity assays were performed as described before (*70*), with minor modification. Cultured NRVMs were lysed on ice in cytosolic extraction buffer (50 mM Tris-HCl pH 7.5, 250 mM Sucrose, 5 mM MgCl2, 0.5 mM EDTA, 1 mM DTT, and 1mM ATP). The cell lysates or myocardium homogenates were plunged through a 29G gauge needle 8 times with an insulin syringe before centrifugation at 4°C for 30 min (15,000 × g). Protein concentration was determined with BCA reagents and samples were diluted with 4× native gel loading buffer (200 mM Tris-HCl, pH 6.8, 60% [v/v] glycerol, 0.05% [w/v] bromophenol blue). Each well of a 4.0% native polyacrylamide gel was loaded with 40 μg of sample and the gel electrophoresis was performed under 15mA constant current for 8 hours. After electrophoresis, the gel was incubated in developing buffer (50 mM Tris pH 7.5, 150 mM NaCl, 5 mM MgCl2, 1 mM ATP) containing 50 μM Suc-LLVY-AMC at 37 °C for 30 min and visualized in a UV trans-illuminator with a wavelength of 365 nm. Immediately after the in-gel peptidase activity assay, the proteins from the native gel were transferred to a PVDF membrane under 250 mA constant current for 90 minutes and immunodetected with anti-Psmb5 antibodies.

### Fluorescence confocal microscopy

Microscopy was performed as described (*19*). Ventricular myocardium from GFPdgn tg mice was fixed with 3.8% paraformaldehyde and processed for obtaining cryosections. The myocardial sections (6 μm) were stained with Alexa Fluor 568-conjuated phalloidin (Invitrogen) to reveal F-actin and identify cardiomyocytes. GFPdgn direct fluorescence (green) and the stained F-actin (red) were visualized and imaged using a Leica TCS SP8 confocal microscope (Leica Microsystems Inc., Buffalo Grove, IL).

### Statistical analysis

Data were statistically analyzed using GraphPad Prism 6 (GraphPad Software, Inc., La Jolla, CA). Western blot and PCR densitometry data are presented as scatter dot plots with mean ± SD superimposed. Differences between two groups were evaluated using two-tailed unpaired Student’s t test. When difference among 3 or more groups was evaluated, one-way analysis of variance (ANOVA) or when appropriate, 2-way ANOVA, followed by Tukey’s multiple comparisons test or Dunnett’s multiple comparisons test were used. Serial echocardiography data were examined using two-way repeated measures ANOVA followed by Bonferroni’s test, to assess interaction of time and drug intervention. For data set with unequal variances, two-tailed unpaired t-test with Welch’s correction was also used and is indicated in figure legends.

## SUPPLEMENTARY MATERIALS

Fig. S1. Verification of the PDE1A antibody.

Fig. S2. Representative images of M mode echocardiograms taken at the completion of 4 weeks of treatment.

Fig. S3. Protein levels of SS fraction of CryAB in NTG and R120G mouse hearts.

Table S1. Echocardiographic measurements at 4 month of Age

Table S2. Echocardiographic measurements at the end of 4 week treatment

## Acknowledgement

We thank Ms. Andrea Jahn and Ms. Megan Lewno for their excellent assistance in transgenic mouse colonies maintenance and genotype determination. We thank the Physiology Core of the Division of Basic Biomedical Sciences for assistance with echocardiography.

## Author contributions

Designing research studies (XW, HZ), conducting experiments (HZ, BP), acquiring data (HZ, BP), analyzing data (XW, HZ, ALG), providing reagents (MDR, ALG), and writing or revising the manuscript (HZ, XW, ALD, MDR, BP).

## Data and materials availability

All original data are available upon request. The transgenic mouse models and recombinant adenoviruses used in this study can be provided to qualified investigators through an MTA.

**This preprint is not peer-reviewed.**

