## Supplemental figures and tables for "Duo-activation of PKA and PKG by PDE1 inhibition facilitates proteasomal degradation of misfolded proteins and protects against proteinopathy"

Fig. S1. Verification of the PDE1A antibody.

Fig. S2. Representative images of M mode echocardiograms taken at the completion of 4 weeks of treatment.

Fig. S3. Protein levels of SS fraction of CryAB in NTG and R120G mouse hearts.

Table S1. Echocardiographic measurements at 4 month of Age

Table S2. Echocardiographic measurements at the end of 4 week treatment

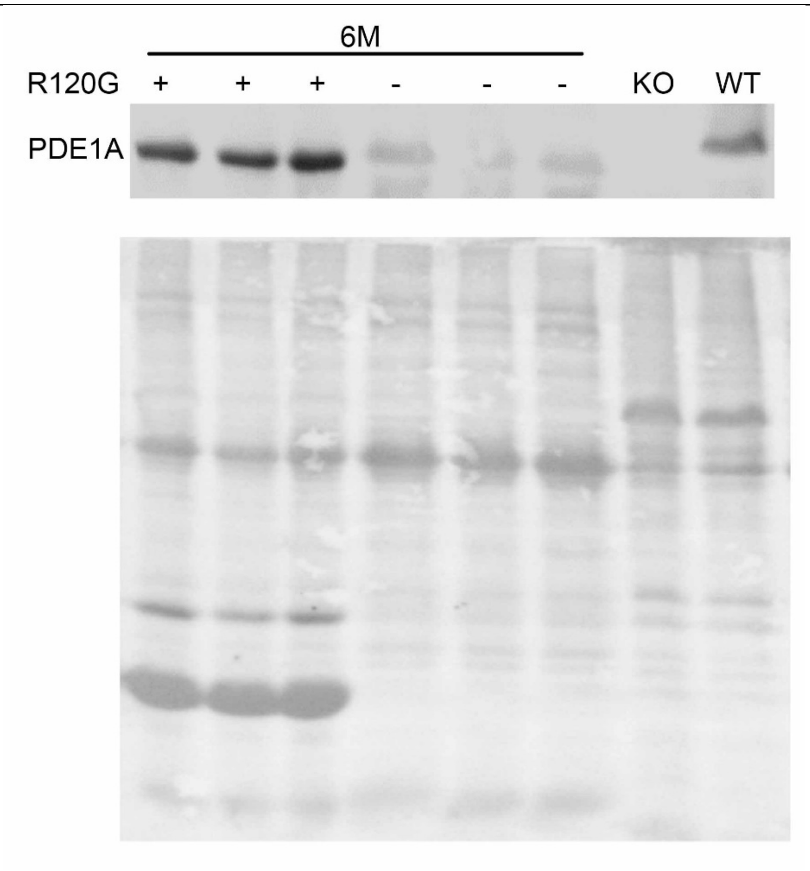


**Fig. S1. Verification of the PDE1A antibody**. Total protein extracts from the brain tissues of wild type (WT) and homozygous PDE1A germline knockout (KO) mice were used for the positive and negative controls in the western blot analyses for PDE1A in the ventricular myocardial tissue from 6 months old CryAB$^{R120G}$ tg and non-tg mice. Note that PDE1A expressed in brain is known to have a greater molecular weight than that expressed in the heart. PDE1A western blot image is shown in the upper panel and the lower panel is the stain-free in-gel labeled total protein image to show the loading.

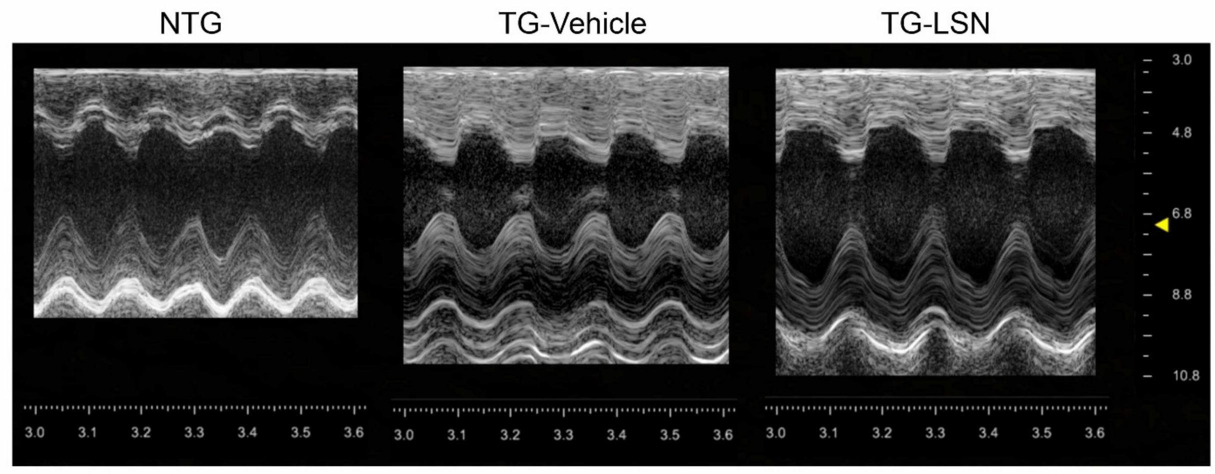


**Fig. S2.** Representative images of M mode echocardiograms taken from sex and age matched non-transgenic (NTG) and CryAB$^{R120G}$ tg mice at the completion of 4 weeks of treatment with the PDE1 inhibitor IC86430 (TG-LSN) or vehicle (TG-Vehicle).

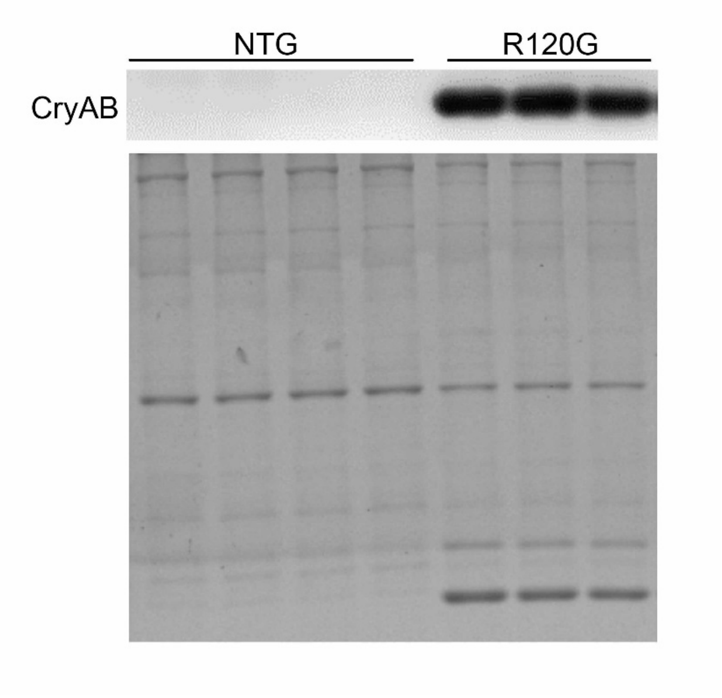


**Fig. S3. Western blot analysis for CryAB levels in the SS fraction of NTG and R120G mouse hearts.** The SS fractions were extracted from ventricular myocardial tissues of ~5 month old non-tg mice and CryAB$^{R120G}$ tg mice as described in the main text. The in-lane total protein image collected with the stain-free technology (bottom) is used as the loading control.

**Table S1.** Echocardiographic measurements at 4 month of age immediately before IC86430 treatment

| | **NTG** (n=8) | **TG-Vehicle** (n=13) | **TG- IC86430** (n=11) |
|---|---|---|---|
| **Body Weight (BW, g)** | 29.4±5.1 | 29.4±5.9 | 27.6±5.5 |
| **Heart Rate (bpm)** | 458±21 | 341±40[§] | 373±49[‡] |
| **CO (ml/min)** | 21.1±3.3 | 13.8±2.5[§] | 14.4±2.5[§] |
| **SV (µl)** | 45.6±6.4 | 36.84±8.3 | 38.7±5.8 |
| **LVVd ($mm^3$)** | 77.41±12.2 | 55.10±10.42[§] | 57.14±7.65[‡] |
| **LVVs ($mm^3$)** | 32.48±10.09 | 18.91±6.23[†] | 18.70±5.00[‡] |
| **LV EF (%)** | 64.40±6.87 | 66.68±7.52 | 66.89±9.72 |
| **LV FS (%)** | 34.67±4.89 | 36.49±6.17 | 37.12±6.90 |
| **LVAWd (mm)** | 0.76±0.121 | 1.20±0.10[§] | 1.15±0.11[§] |
| **LVAWs (mm)** | 1.04±0.10 | 1.48±0.15[§] | 1.50±0.10[§] |
| **LVIDd (mm)** | 4.14±0.29 | 3.61±0.26[†] | 3.63±0.18[†] |
| **LVIDs (mm)** | 2.90±0.35 | 2.35±0.36[†] | 2.39±0.27[†] |
| **LVPWd (mm)** | 0.76±0.13 | 1.17±0.14[§] | 1.11±0.11[§] |
| **LVPWs (mm)** | 1.09±0.08 | 1.49±0.16[§] | 1.50±0.10[§] |
| **LV Mass (mg)** | 95.0±20.6 | 150.1±30.3[†] | 140.2±21.2[§] |
| **LV Mass/BW (mg/g)** | 3.23±0.46 | 4.95±0.99[‡] | 5.19±0.94[§] |

LVIDd, end-diastolic left ventricular (LV) internal diameter; LVIDs, end-systolic LVID, end-diastolic LV posterior wall thickness; LVPWs, end-systolic LVPW; LVAWd, end-diastolic anterior wall thickness; LVAWs, end-systolic LVAW; FS, fractional shortening; EF, ejection fraction; LVVd, end-diastolic LV volume; LVVs, end-systolic LVV; SV, stroke volume; CO, cardiac output. For all parameters, there is no statistically significant difference between the TG-Vehicle group and the TG- IC86430 group immediately before IC86430 treatment.

*p<0.05, †p<0.01, ‡p<0.001, §p<0.0001 vs. NTG; one-way ANOVA followed by Tukey's multiple comparison tests

**Table S2.** Echocardiographic measurements at the end of 4 week treatment

| | **NTG** **(n=8)** | **TG-Vehicle** **(n=8)** | **TG- IC86430** **(n=11)** |
|---|---|---|---|
| **Body Weight (BW, g)** | 30.5±4.2 | 29.8±5.0 | 29.7±4.3 |
| **Heart Rate (bpm)** | 462±53 | 361±59$^{\ddagger}$ | 344±50$^{\S}$ |
| **CO (ml/min)** | 21.5±4.1 | 12.5±0.9$^{\#}$ | 16.1±3.2$^{\ddagger}$* |
| **SV (µl)** | 46.5±10.4 | 36.3±4.9$^{\parallel}$ | 46.5±7.1* |
| **LVVd (mm$^3$)** | 73.54±12.5 | 57.83±4.99$^{\parallel}$ | 72.63±13.55* |
| **LVVs (mm$^3$)** | 27.08±4.39 | 21.59±4.61 | 28.11±8.61 |
| **LV EF (%)** | 59.76±4.19 | 62.77±7.19 | 62.87±6.63 |
| **LV FS (%)** | 31.42±2.86 | 33.58±5.72 | 33.82±4.73 |
| **LVAWd (mm)** | 0.75±0.06 | 1.20±0.07$^{\#}$ | 1.07±0.06$^{\#\dagger}$ |
| **LVAWs (mm)** | 1.05±0.05 | 1.49±0.08$^{\#}$ | 1.35±0.08$^{\#\dagger}$ |
| **LVIDd (mm)** | 3.95±0.31 | 3.71±0.09 | 4.07±0.28* |
| **LVIDs (mm)** | 2.75±0.25 | 2.53±0.27 | 2.89±0.40 |
| **LVPWd (mm)** | 0.78±0.08 | 1.18±0.11$^{\#}$ | 1.07±0.08$^{\#}$* |
| **LVPWs (mm)** | 1.05±0.07 | 1.51±0.11$^{\#}$ | 1.36±0.10$^{\#}$* |

LVIDd, end-diastolic left ventricular (LV) internal diameter; LVIDs, end-systolic LVID, end-diastolic LV posterior wall thickness; LVPWs, end-systolic LVPW; LVAWd, end-diastolic anterior wall thickness; LVAWs, end-systolic LVAW; FS, fractional shortening; EF, ejection fraction; LVVd, end-diastolic LV volume; LVVs, end-systolic LVV; SV, stroke volume; CO, cardiac output.

*$p$<0.05, $^{\dagger}p$<0.01 vs.TG-Vehicle,

$^{\parallel}p$<0.05, $^{\ddagger}p$<0.01, $^{\S}p$<0.005, #$p$<0.001 vs. NTG, one-way ANOVA followed by Tukey's multiple comparisons test.
